# Comparing miRNA structure of mirtrons and non-mirtrons

**DOI:** 10.1101/218701

**Authors:** Igor I. Titov, Pavel S. Vorozheykin

## Abstract

**Background:** MicroRNAs proceeds through the different canonical and non-canonical pathways; the most frequent of the non-canonical ones is the splicing-dependent biogenesis of mirtrons. We compare the mirtrons and non-mirtrons of human and mouse to explore how their maturation appears in the precursor structure around the miRNA.

**Results:** We found the coherence of the overhang lengths what indicates the dependence between the cleavage sites. To explain this dependence we suggest the 2-lever model of the Dicer structure that couples the imprecisions in Drosha and Dicer. Considering the secondary structure of all animal pre-miRNAs we confirmed that single-stranded nucleotides tend to be located near the miRNA boundaries and in its center and are characterized by a higher mutation rate. The 5′ end of the canonical 5′ miRNA approaches the nearest single-stranded nucleotides what suggests the extension of the loop-counting rule from the Dicer to the Drosha cleavage site. A typical structure of the annotated mirtron pre-miRNAs differs from the canonical pre-miRNA structure and possesses the 1- and 2nt hanging ends at the hairpin base. Together with the excessive variability of the mirtron Dicer cleavage site (that could be partially explained by guanine at its ends inherited from splicing) this is one more evidence for the 2-lever model. In contrast with the canonical miRNAs the mirtrons have higher snp densities and their pre-miRNAs are inversely associated with diseases. Therefore we supported the view that mirtrons are under positive selection while canonical miRNAs are under negative one and we suggested that mirtrons are an intrinsic source of silencing variability which produces the disease-promoting variants. Finally, we considered the interference of the pre-miRNA structure and the U2snRNA:pre-mRNA basepairing. We analyzed the location of the branchpoints and found that mirtron structure tends to expose the branchpoint site what suggests that the mirtrons can readily evolve from occasional hairpins in the immediate neighbourhood of the 3′ splice site.

**Conclusion:** The miRNA biogenesis manifests itself in the footprints of the secondary structure. Close inspection of these structural properties can help to uncover new pathways of miRNA biogenesis and to refine the known miRNA data, in particular, new non-canonical miRNAs may be predicted or the known miRNAs can be re-classified.

## Background

Canonical pathway of animal miRNA begins with transcription. RNA polymerase II (or polymerase III for some miRNAs) creates long primary transcript (pri-miRNA) which contains one or more hairpins (pre-miRNAs), poly(A) tail and 7-methylguanosine cap [1, 2]. MiRNA genes are dispersed in various genomic locations (intronic, exonic or intergenic regions) and can be transcribed independently or as a part of other host genes [3–5]. A cluster brings together miRNAs with inter-miRNA distance up to 10kb and can form a polycistronic transcriptional unit (for example, mir-100/let-7/mir-125 and mir-71/mir-2 clusters) [6–8]. MiRNAs can be located in both DNA strands (for example, hsa-miR-3120 and hsa-miR-214, dme-miR-iab-4): although these miRNAs are close to each other, they can be regulated post-transcriptionally either united or independent [9, 10].

After the transcription, animal pri-miRNAs are cleaved by the Microprocessor complex of the RNase III enzyme Drosha and its co-factor DGCR8 [3]. The complex releases pre-miRNA hairpin by cropping the stem-loop [11–13]. This step can be regulated by a variety of ways: in some of them proteins are recruited to protein-protein interactions, in others the pri-miRNA primary and secondary structures are involved in RNA-protein or RNA-RNA bindings.

Further, the Drosha product is moved from the nucleus to the cytoplasm by the protein Exportin-5 (EXP5) and the cofactor Ran-GTP [14]. Some other proteins (for example, XPO1) can transport non-canonical pre-miRNAs [15]. EXP5 does not only transfer the precursors, but also prevents them from degradation [16].

In the cytoplasm, the pre-miRNA must be cleaved by RNase III enzyme Dicer near the terminal loop. The cleavage releases a double stranded miRNA duplex with typical 2nt 3′ overhangs [17]. Usually, animal Dicer contains the following domains: helicase, PAZ, dsRNA binding and two RNase III (A and B) domains [18]. Each of these domains are involved in the miRNA maturation process. The helicase domain promotes the pre-miRNA recognition by interacting with the terminal loop and facilitates the processing [19]. The PAZ domain identifies the precursor’s termini and binds to them. Each of the two RNase III domains cuts one of the two pre-miRNA strands and releases the miRNA duplex from the terminal loop [20, 21].

After the Dicer has produced the miRNA duplex, a miRNAs-induced silencing complex (miRISC) is formed and targets mRNAs [22–24] or non-coding RNAs [25–27].

In addition to the canonical miRNA biogenesis described above, another pathways can generate miRNAs in a Drosha- and/or Dicer-independent manner [28, 29]. Most of the non-canonical miRNAs are mirtrons which bypass the Drosha cleavage step and are derived through the mRNA splicing, the lariat debranching and refolding into a canonical-like stem-loop structure [30–34]. If this stem-loop contains extra-nucleotides at 5′ or 3′ ends (the so-called “tailed” mirtron), they are trimmed by exonucleases to gain an appropriate structure for the exportin complex. From this time on, the processing goes on the canonical pathway. At present, the hundreds of mirtrons have been found [32], however the features of non-canonical maturation are still poorly studied in contrast with the canonical one.

Mammalian mir-1225 and mir-1228, initially predicted as mirtrons, are actually splicing-independent [35]. Moreover, biogenesis of these miRNAs does not require the most of the canonical components (DGCR8, Dicer, Exportin-5 or Ago2) but still involves Drosha [36, 37]. This class of miRNAs, termed “simtrons” (splicing-independent mirtron-like miRNAs), reveals a new pathway of small regulatory RNA production [36, 37]. Another Dicer-independent pathway is observed for the mir-451 family, this pathway involves the catalytic activity of the Ago2 protein [38–40]. Drosha generates pre-mir-451 with ~18-nt stem which is too short to be processed by Dicer. Therefore the pre-miRNAs are processed by Ago2 which cleaves the hairpin in the middle of its 3' strand and yields a ~30nt long RNA product [38–40]. Then poly(A)-specific ribonuclease PARN trims the 3′ RNA end to release the mature 5′ miRNA [41].

On each step of miRNA maturation, the biogenesis leaves its footprints as the specific pre-miRNA and miRNA features. The canonical miRNAs are usually located near the terminal loop of the pre-miRNA hairpin. Simultaneously some non-canonical miRNAs (e.g. originated from simtrons mir-1225 and mir-1228) are distant from the terminal loop. The mir-451 family can be processed by the canonical pathway as well as by the non-canonical one in which Drosha produces a short hairpin with the miRNA that overlaps within the terminal loop. Based on the pri-/pre-miRNA structural properties, new non-canonical miRNAs may be predicted or the known miRNAs can be re-classified. Also, close inspection of these characteristics can help to uncover the new pathways of the miRNA biogenesis and to discover the errors in annotated miRNA data.

It is commonly believed [42] that miRNA genes have been either evolved from random hairpins in intergenic regions or in intronic regions of protein-coding genes or duplicated the miRNA genes and the transposable elements (e.g. the fraction of the human TE-derived miRNAs in miRBase had been constantly growing [43]). Most of new miRNAs disappeared over time while the survived ones adapted and then came under purifying selection like the old miRNAs until they could start an another cycle of adaptive-conservative evolution in other tissues [44]. Many of these new miRNAs are mirtrons, they have been often evolved in clade- and species-specific ways and more quickly than the canonical miRNAs [45, 46].

In this paper we consider the structural properties of the animal miRNAs and compare mirtrons with non-mirtrons, most of the latter are the canonical miRNAs. First, we study the miRNA pair layout which shows itself in overhang lengths. Second, we investigate the distances from the miRNA ends to the nearest single-stranded nucleotide. Then we inspect the loop positional frequencies in the miRNA and its flanks and correlate these frequencies with the mutation rate. Next, we study SNP density in miRNA and its flanks. Finally, we consider how the branchpoints are located within the mirtron pre-miRNAs.

Our observations support the current view that RNA secondary structure plays a crutial role in miRNA maturation and exemplify how the biogenesis peculiarities become apparent in this structure. A lot of miRNA/pre-miRNA prediction methods use the RNA secondary structure [47–54], so our results can be useful for further improvement of the existing methods. The excessive SNP density and the branchpoint locations within mirtron precursors demonstrate that mirtrons represent new miRNAs which could be easily recruited from introns. The further inspection of the mirtron branchpoints can help in better understanding the role of the secondary structure in splicing.

## Methods

The sequences and structures of the pre-miRNAs and miRNAs were downloaded from the miRBase database (release 21.0) [55]. There are 15 731 unique animal pre-miRNAs which contain 22 603 experimentally validated mature miRNAs approximately equally in both arms of the precursors. We excluded few pre-miRNAs with non-canonical nucleotides and with more than two annotated miRNAs.

We selected those mirtrons which are simultaneously presented both in miRBase-21.0 and in the paper [56]. These data contain 464 human and mouse mirtron pre-miRNAs, while a number of non-mirtron human and mouse pre-miRNAs is 2438.

The revisited miRNA sequences were taken from the miRBase-21.0 whose identifiers are simultaneously presented in [57]. This set contains more than one thousand animal pre-miRNAs. We used the SNP database miRNASNP-2.0 (based on miRBase-19.0 and dbSNP137) [58] to calculate the SNP densities in human miRNA genes and their flanks. The dataset contains both common (minor allele frequency > 0.01) and rare SNP variants. The SNP density was defined as: *N*_*Snp*_ x 1000/L, where *N*_*snp*_ was the number of the SNPs in the RNA region, *L* was the length of the region (seed, miRNA excluding seed, pre-miRNA excluding miRNA). SNP occurrence per sequence for disease and non-disease human pre-miRNAs was calculated as in [59] and was based on miRNA associated diseases from [60] and on miRNASNP-2.0 [58].

Branchpoint data of human and mouse introns were taken from the supplemental materials of the paper [61].

Data of the animal nucleotide substitutions were taken from [62].

The unpaired nucleotide frequency (UNF) for each RNA position was calculated as the portion of miRNAs that had a single-stranded nucleotide at the position.

The distance between miRNA end and its nearest single-stranded nucleotide was calculated as the minimal number of the nucleotides between the miRNA boundary and the single-stranded region in the same miRNA strand.

## Results and Discussion

### Overhang lengths

The diversity of the miRNA cleavage site leads to overhang variety, therefore they can shed new light on the nature and mechanism of the cleavage process. The overhangs are the miRNA ends hanging from its miRNA duplex thus reflecting the miRNAs disposition. Both Dicer and Drosha cut the miRNA precursor, especially the 3' variable miRNA ends, in a number of neighbouring positions and can form other than canonical 2nt overhangs. Each cleavage variant produces its own version of the miRNA duplex and in some miRBase records the variant with 2nt overhangs is not the most observable.

The Dicer and Drosha interactions with pre-miRNA, especially sensitivity of their RNase domains (RIIIA and RIIIB) to RNA sequence, defines the miRNA ends. These domains prefer to generate the U-ended miRNAs as the main fraction [63]. The G-ended miRNAs rarely occur and the G-avoiding generates the atypical 1nt and 3nt overhangs [64]. For the “homogeneous” cleavage (as it was defined by [63]) the overhang shortening appears as a 1nt context shift at the 3′ miRNA ends (compare figure 2A and figure 2С of the second-most frequent miRNA fraction in [63]). Sometimes these shortened miRNAs are found in miRBase in another species: compare, for example, tgu-let-7b, aca-let-7b and mml-let-7b miRNAs (Additional file 1). Occurrence of the A/G at the neighbourhood of miRNA ends could lead to the heterogenious cleavage, i.e. to levelling of the miRNA fractions (figure 2 from [63]). Based on the works of Starega-Roslan et.al. [63–66], one can conclude that the overhang lengths are sequence-dependent; they are also structure-dependent as it was observed in [67] where the sliding (bulge) loops induced cleavage heterogeneity.

Since not only the sequence but also the structure of the pre-miRNA can trigger this cleavage heterogeneity, we measure the overhang lengths by a number of excessive nucleotides beyond the closing pair of the miRNA duplex regardless of its structural state, single-stranded or double-stranded, rather than a length of hanging end.

To estimate the occurrence of the atypical overhangs we study the overhang lengths, considering the mirtrons as a separate miRNA class. Unlike the canonical miRNAs, the mirtrons use splicing to bypass Drosha cleavage. The mirtron database consists mainly of human and mouse mirtrons [56], therefore we consider four miRNA sets: animal miRNAs (1), animal miRNAs without human and mouse ones (2), human and mouse non-mirtrons (3) and mirtrons (4). The third set contains only few numbers of known non-canonical miRNAs which could not significantly influence on.

Figures 1A–1C show the overhang length distributions of the miRNA duplexes from three of four sets. The data on figure 1A includes the canonical miRNAs, Drosha-independent mirtrons and a small number of other non-canonical miRNAs, e.g. Dicer-independent mir-451 family [29], simtrons [37], etc. As we see, figure 1A does not significantly differ from the figure 1B which represents the distributions of human and mouse non-mirtrons. The data for all animal miRNAs are also similar to figure 1A and figure 1B and therefore are not shown here. Figure 1C displays the corresponding distributions of the most abundant non-canonical class, mirtrons.

**Figure 1.**
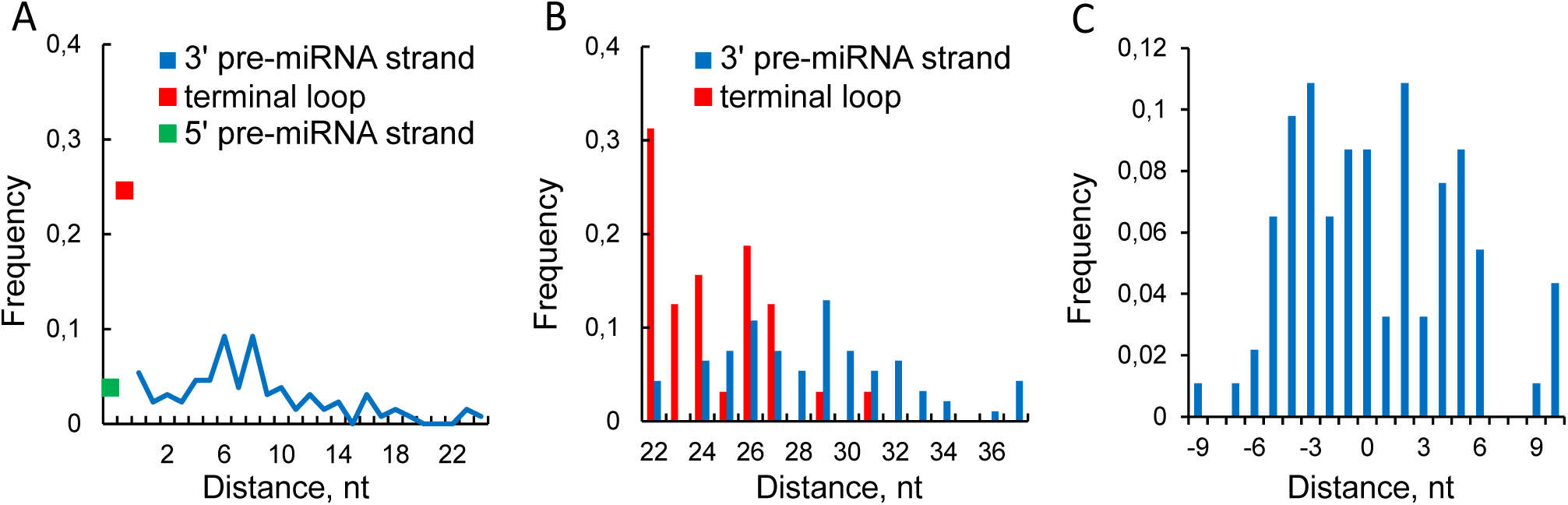
The overhang lengths of miRNA duplexes. The frequency of the overhang lengths of miRNA duplexes: animal miRNAs without human/mouse ones (A), human and mouse non-mirtrons (B) and mirtrons(C). The overhang lengths occurrence of both cleavage sites for animal miRNA duplexes; in each quarter-square the miRBase pre-miRNA structure which leads to the corresponding overhang types is schematically shown (D). Negative values correspond to an atypical 5' overhangs. The long overhangs on the panel D correspond to structure prediction errors and are described further in the text. These overhangs are not shown on the panels A and C. In mirtron case the splicing overhangs are considered instead of the Drosha ones.

The canonical overhang lengths for both cleavage sites are well-known to be equal to 2nt, what is actually observed on figures 1A and 1B where all the length distributions peak at 2nt. The overhang distributions of the Dicer and Drosha cleavage sites are similar and asymmetrical (figures 1A and 1B). The overhangs are more readily shortening what could be explained either by more frequent exosome cutting of the 3′ miRNA end than of the 5′ one [68] and/or by the 3′ cleavage site shifting inside the duplex.

The Drosha overhang distribution closely matches the Dicer’s one (figure 1B) what supports the observations that Drosha and Dicer process the canonical miRNAs in a similar manner. In the mirtron case the splicing replaces the Drosha step, but the overhang statistics are surprisingly changed for both cleavage sites (figures 1B and 1C). First, splicing overhang distribution is shifted relatively to the Dicer distribution (figure 1C) what can reflect the exonuclease trimming of 5′-tailed mirtrons which are most frequently observed [56]. Second, although the same complex cleaves both mirtrons and non-mirtrons at the Dicer site, the overhang distributions differ (figures 1B and 1C): mirtrons distribution blurs what suggests lower Dicer cleavage precision, presumably due to the dependence of Dicer result on the output of splicing and further exonuclease editing.

The introns usually have sequence conservations at the 5′ end (GU) and at the 3′ end (AG). While the majority of mirtrons are tailed and lose one of the intron ends during the exosome cutting, the remaining end can contribute to the Dicer cleavage heterogeneity as it follows from the figure 2 in [63].

**Figure 2.**
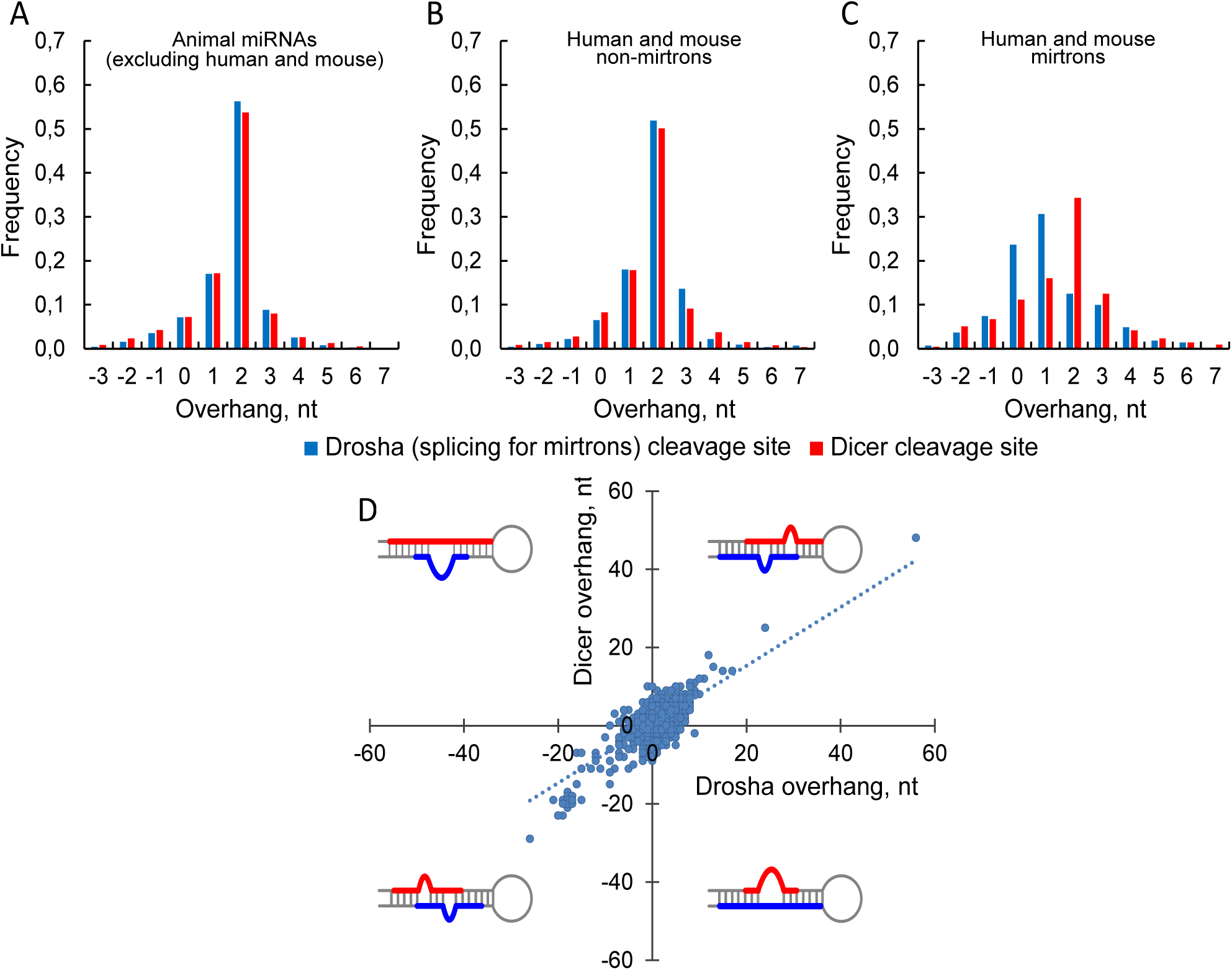
Distance between miRNA end and its nearest single-stranded nucleotide in the same miRNA strand. Considered are only those miRNA ends in which their terminal nucleotide is double-stranded. The frequencies of the 5′ miRNA ends are shown on panels A and C. The frequencies of the 3′ miRNA ends are shown on panels B and D. The data are presented for 5′ and 3′ miRNA sequences separately: for animal miRNAs excluding human and mouse ones (A and B) and for the human and mouse mirtrons and non-mirtrons (C and D). The positive values correspond to the distances to the nearest single-stranded nucleotide outside the miRNA. The negative values are the numbers of nucleotides that must be cut off from the miRNA to reach the nearest loop in the miRNA. The distance 0 is observed for those miRNA ends that are exactly at the boundary of the single-stranded region. The distance frequencies for all animal miRNAs (not shown) are the sum of results for all datasets and are only slightly different from the observations A and B.

If the Dicer heterogeneity for mirtrons is caused not only by guanine at their ends, but also by a heterogeneity of the overhang lengths of splice site, such a dependence should appear in a coordinated variation of Dicer and Drosha overhangs of canonical miRNAs. This contradicts to the fact that Drosha and Dicer cleave independently and to clear this discrepancy up we plotted the dependence of the overhang lengths for both cleavage sites of animal miRNAs (figure 1D). Indeed, we observe significant linear dependence (ρ=0.338, P = 2.42×10^-183^, Spearman’s rank correlation test) of the Dicer and Drosha overhang lengths (figure 1D). This dependence arises due to the several reasons. First, the way the pre-miRNA structure is formed, it excludes big bulge loops within miRNA duplex and, consequently, the pairs of long overhangs of the opposite sign. Second, the pairs of long overhangs of the same sign are observed due to the incorrect prediction of terminal loop (Additional file 2) or to the presence of false miRNAs in the miRBase. And the last possibility is the guanine avoiding at the first position of 5′ end of 3′ miRNA [63, 64]. To exclude these three reasons, we further considered only the canonical overhangs and the overhangs with minimal 1nt deviations from them. The first two reasons disappeared due to the near-canonical overhang lengths, and the last reason (at least as a main factor) was excluded after comparing guanine frequencies in three neighbouring positions at the miRNA boundary (see table 1 in Additional file 3). The remained duplexes with these weakly varying overhang lengths compose the most part (67.7%) of all duplexes and their overhangs still significantly correlate (ρ=0.139, P = 2.2×10^−21^, Spearman’s rank correlation test).

The overhang lengths interdependence can be also verified in biochemical studies (for example [69]). Unfortunately, their paper does not provide the data on joint occurrence of miRNA/miRNA* and therefore can not be used to test our hypothesis.

To understand the nature of this correlation, as the null hypothesis we considered the 2-parametric model of independent overhang lengths and fitted the length frequencies (table 2, Additional file 3). For the both (Dicer and Drosha) cleavage sites the long (short) overhangs are observed about 7 (4) times less often than the canonical ones (table 3, Additional file 3) as it was already seen in figure 1A and figure 1B This model describes well all length frequencies except for the1nt/1nt and 3nt/3nt length pairs which are observed twice as often as expected. So, these pairs are what induce the length correlation. Moreover, this correlation is robust to definition of the overhang length (Additional file 4).

We suggest that this correlation reflects the organization of the pre-miRNA/Dicer complex. The pre-miRNA/Dicer complex consists of the sub-units that move as a whole, what appears as a collective movements of large scale around a hinges. The PAZ domain is the main moving domain for the Dicer [70] this domain adapts to the pre-miRNA ends. We speculate that among all possible collective movements pre-miRNA/Dicer complex undergoes smaller scale movements of 2-lever type which are responsible for the coherence trend of the overhang lengths (Additional file 5). The tips of two levers bind to the pre-miRNA ends in the PAZ domain. The another two tips of the levers are located in the RNase IIIA and RIIIB determining the distance between the cleavage sites. As a result, the tips of the levers can close in and move away in concert what leads to such a number of states of the cleavage complex that the Dicer overhang tends to vary cooperatively with the Drosha one.

Unfolding this 2-lever model we note that these levers should differ in their rigidity. One lever (associated with Dicer RNase IIIA) is connected with the 5′ ends of the miRNAs and tightly bound to RNA which, as well as the connector helix, ensures the lever rigidity and manifests itself in less variability of the 5' end of the 3' miRNA and in G-avoiding on both 5′ ends. In contrast, the other lever (associated with Dicer RNase IIIB) is soft and its free movement forms the cleavage variability and the variety of the overhang lengths.

Another evidence in favor of the lever mechanism is the more pronounced heterogeneity of the Dicer cleavage site for mirtrons in which the overhang lengths of the splicing site are more variable and the ends are often freely hanging (see next two sections of this paper) thus forming different spatial distances between each other.

### miRNA end distance to the nearest single-stranded region

The pre-miRNA secondary structure, as well as the nucleotide sequence, can influence the miRNA boundaries. The Dicer cleavage depends either on the stem size or the terminal loop [71] and its precision (i.e., the fraction of the most probable miRNA) also fulfills the so-called “loop-counting rule”: Dicer cleaves precisely at 2nt distance to any upstream loop, in other cases Dicer produces variable 5′ end of the 3′ miRNA [69].

Gu and co-authors were focused on the Dicer cleavage site. We inspect how the loop-counting rule makes itself evident in the distance between miRNA end and the single-stranded regions and answer the question “Is there something like the loop-counting rule for the Drosha cleavage site?”. Expecting structural difference between canonical and non-canonical pre-miRNAs we explore separately mirtrons and non-mirtrons.

As one can see on figure 2A the loop-counting rule appears as a pronounced peak at 2nt which contains approximately 40% of animal miRNAs (excluding human and mouse ones). The remaining cases are partially referred to the incorrect prediction of the pre-miRNA secondary structures (Additional file 2) and to the fact that the structure is locally unstable in the loop neighbourhood. The blue peak (5′ end of the 5′ miRNA) is even more pronounced, this miRNA end is located immediately before the single-stranded region (figure 2A). This suggests that the loop-counting rule for the Drosha cleavage site exists as well.

Figure 2B shows the distance frequencies to the 3′ miRNA end. The broader distribution on figure 2B comparing with the figure 2A suggests that the cleavage complex for the 3′ miRNA end is less accurate than for the 5′ end as it was already proposed by [63]. Both cleavage sites are processed by Drosha and Dicer RNase domains (RIIIA and RIIIB). The RIIIA domain processes the 3′ miRNA while the RIIIB domain handles the 5′ one. Therefore, the RNA site affects the cleavage precision to a greater extent than the particular RNase domain. Specifically, the 5′ end is controlled by well-defined neighbouring structure (figure 2A) and nucleotide sequence [63]. In contrast, the 3′ end may be determined mostly by the overhang length (or distance between RNase IIIA/IIIB cleavage sites according to the lever model) thus providing the required size of the miRNA.

Some of these animal miRNAs are mirtrons which are though rare but the most abundant non-canonnical miRNAs. To reveal the structural diffirence between canonical and non-canonical miRNAs we consider miRNAs of human and mouse where mirtrons are better identified.

The frequency distributions near the hairpin terminal loop are similar for mirtrons and non-mirtrons (red and pink bars on figure 2C, blue and light blue bars on figure 2D) what reflects the fact that the Dicer processes the both classes in the same way. In contrast, the mirtron and non-mirtron frequency distributions strongly differ for another cleavage site where the different processing complexes cleave the RNA molecules (red and pink bars on figure 2D, blue and light blue bars on figure 2C). More pronounced peaks of mirtrons stems from the observation that their ends at the hairpin base are located at 0-1nt distance from the single-stranded region (figures 2C and 2D). Some of these mirtron pre-miRNAs are not produced immediately by splicing, but are rather derived by further cropping the single-stranded regions by exonucleases.

Thus the pre-miRNA secondary structure is an important factor for precise recognition of both miRNA ends. In particular, the loop-counting rule [69] could be extended to the Drosha cleavage site. The structural signature near the hairpin base of the mirtron pre-miRNAs is so clearly defined that may be useful for their validation.

### The unpaired nucleotide frequency across miRNA

As we have seen above, loop position is an important factor of miRNA processing. Therefore, we consider the unpaired nucleotide frequencies (UNFs) within miRNA and how these frequencies relate to nucleotide substitutions. Going along the secondary structure of miRNA classes we compare mirtron and non-mirtron unpaired nucleotide frequencies (UNFs) within miRNA and its nearest neighbourhood (figures 2C and 2D).

Figure 3A shows that the UNF varies across the miRNA sequence. The miRNA positions can be roughly divided into two groups (figures 3A and 3B). The first group, loop-rare positions (grey region), contains the seed region (positions 2-8) and the additional binding site (positions 13-16) with few nucleotides downstream (positions 17-19). The UNFs of these regions are close because they are typically paired to each other in the opposite strands of the miRNA duplex. The loop-frequent positions (white region) fall into miRNA center and its ends.

**Figure 3.**
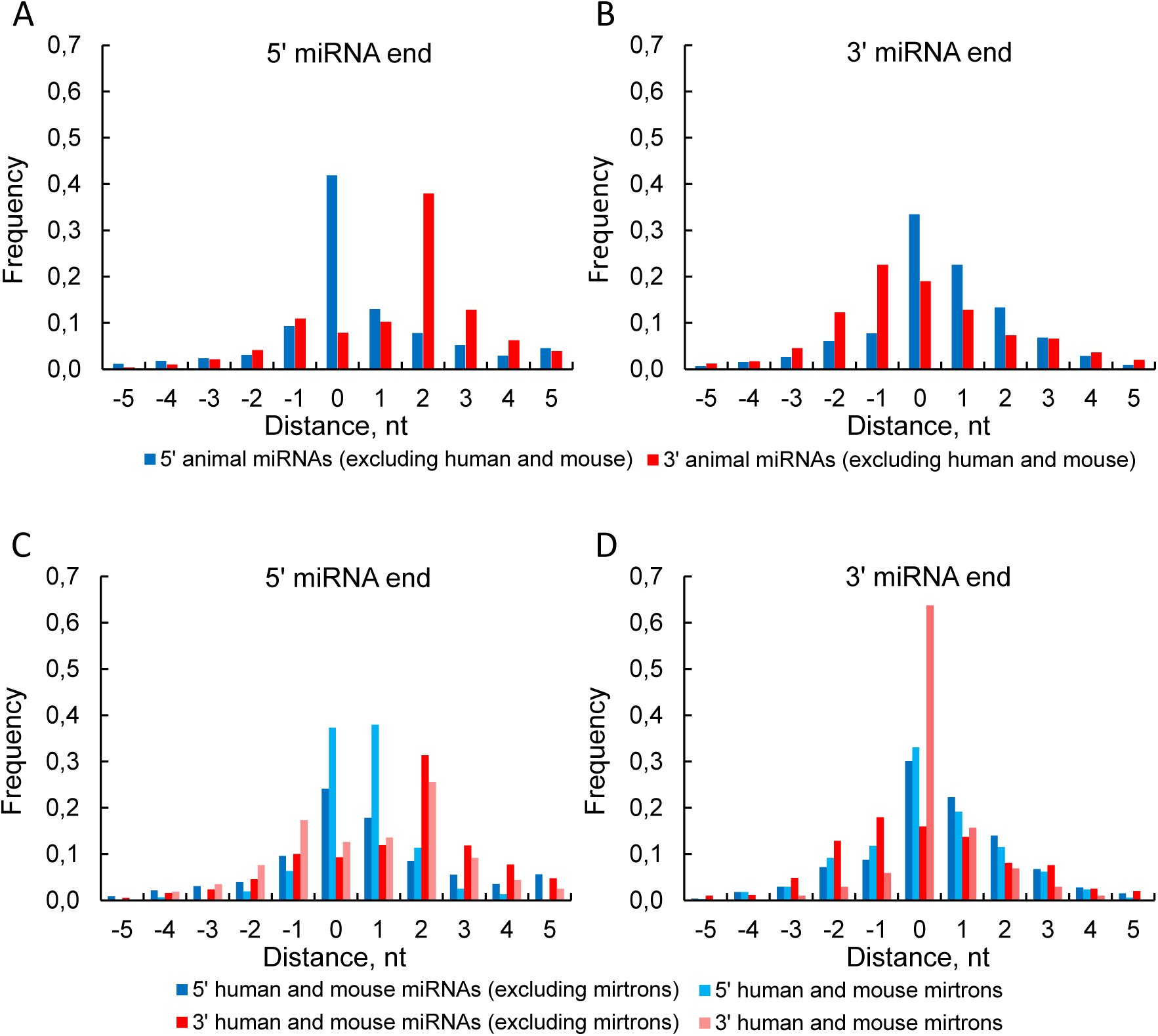
The unpaired nucleotide frequency (UNF) across the miRNA sequence. 5′ end of the miRNA starts from position one. Negative positions correspond to the miRNA flank. The UNF is not shown at the very ends of several long miRNAs. (A) Animal miRNAs. (B) The UNF dependence on the relative rate of nucleotide substitutions in animal miRNAs [62]. The seed points concentrate near the very UNF-axis. Spearman’s rank correlation test was used to estimate the significance of the correlation between the UNF and the rate of nucleotide substitutions (ρ=0.81, P=2.76×10^-6^). (C-D) The UNF profile of 5′ miRNAs and of 3′ miRNAs of human and mouse (mirtrons and non-mirtrons) and of animal excluding human and mouse.

Wheeler reported that the substitution rate reflects the importance of the positions 2-8 and 13-16 [62]. To reveal the relation between secondary structure and mutations within miRNA sequence we plotted the dependence of the UNF on the nucleotide substitution rate taken from [62]. Although the correlation between the UNF and the substitution rate is significant (P=2.76×10^-6^), this dependence is stepwise rather than linear (figure 2B). The step is formed by two groups of positions (grey and white regions) in each of them the dependence does not exist. The seed region (positions 2-8) contains the most conserved positions in the miRNA [62]. Four neighboring positions of the seed and of the additional binding site (positions 9 and 17-19) form the transition from the conserved double-stranded to the more variable single-stranded regions of the miRNA. This agrees with the fact that a RNA base-pairs near a loop use to be partially unwinded. Taking together figures 3A and 3B we conclude that the secondary structure is one of the miRNA evolutionary constraints in the same way as for the majority of structural RNAs where the loops are more variable.

On figures 3A and 3B we observe three variable and, at the same time, single-stranded miRNA regions: the center and the ends. The 5′ miRNA end together with the mismatches at miRNA center are responsible for Ago-protein sorting, as it was shown for some animal species [71]. Figures 3A and 3B (as well as figure 2) represent the miRNA ends effort to be bounded by singe-stranded nucleotides. As a part of this tendency, the left (right) peak on figure 3C (3D) supports again the existence of the loop-counting rule for the Drosha cleavage site. As for mirtrons they have a similar UNF profile over the whole sequence except the positions by the hairpin base (figures 3C and 3D), where the mirtrons are much more single-stranded what characterizes their unique biogenesis. In particular, the first nucleotide of the most 5′ mirtrons (79%) is single stranded and quite often survives after the intron cropping by exonucleases [31]. Its opposite end of the 3′ pre-miRNA is also often single-stranded and uses to be immediately formed by splicing [31].

### SNP density of human miRNA and its neighbourhood

SNPs are the most frequent genetic changes in human genome. The miRNA-related SNPs may affect the miRNA functions and subsequently result in the phenotype changes and diseases so that some miRNAs could appear as the diseases-prediction biomarkers. These SNPs could potentially alter miRNA maturation, silencing machinery, pri-/pre-/miRNA structure, miRNA expression and target binding [72].

The previous papers have produced the controversial values of the SNP densities (red and yellow bars, figure 4A) [73, 74]. Therefore, we recalculated the SNP density using the latest human SNPs in the pre-miRNAs and their flanks. Returning to the keynote of our paper we consider separately the SNP densities of non-mirtrons and mirtrons (blue and green bars, correspondingly, figure 4A).

**Figure 4.**
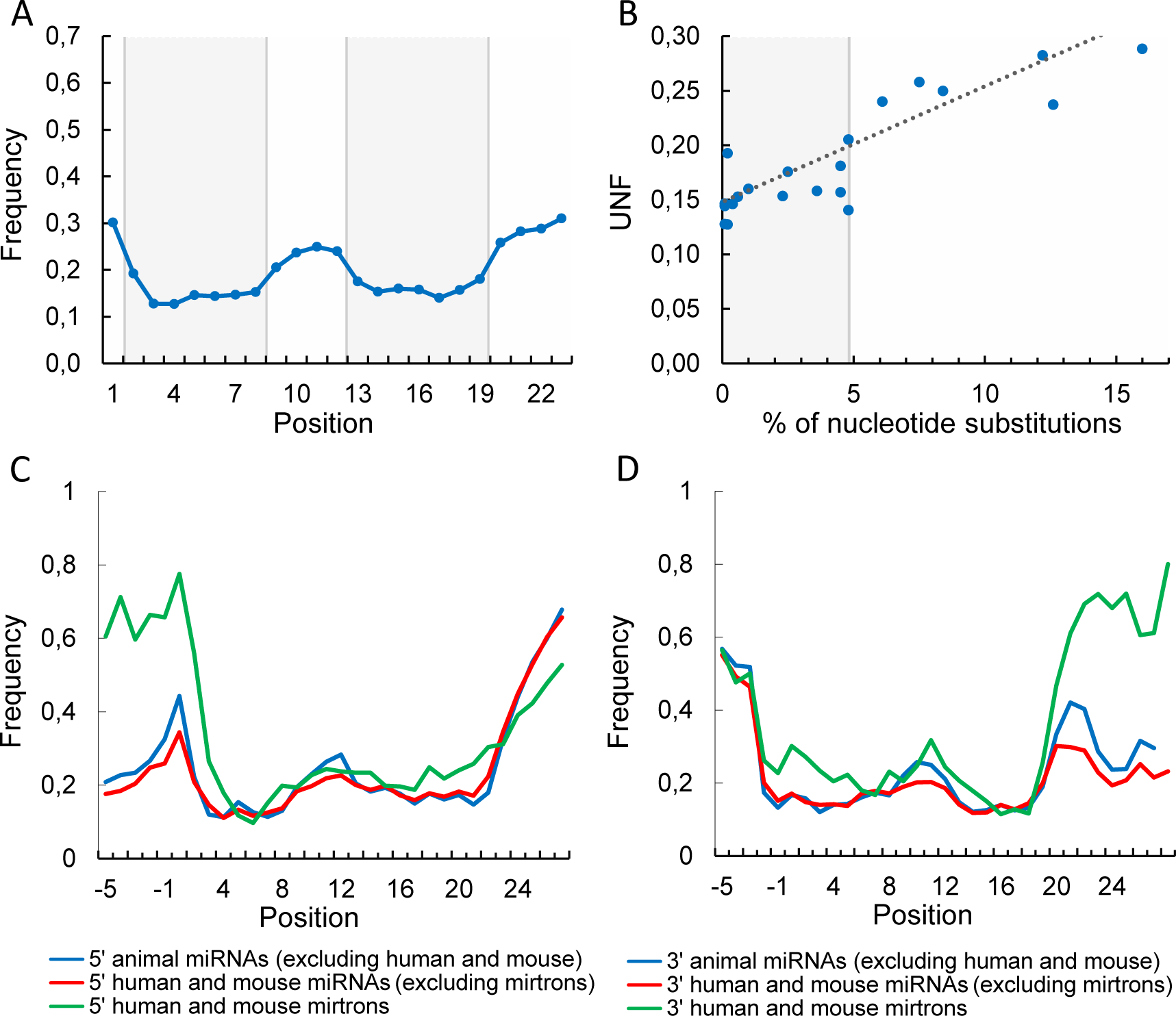
SNP in pre-miRNA and its link with diseases. (A) SNP density in human pre-miRNAs and their flanking regions. Blue (green) bars correspond to human non-mirtrons (mirtrons). The results are based on miRNASNP-2.0 (miRBase 19.0 and dbSNP137) [58]. Red bars were calculated by [73] using miRBase-16.0 and dbSNP132 (miRNASNP-1.0 database). Yellow bars were calculated by [74] using miRBase 18.0 and dbSNP135 (MirSNP database). SNP density is shown separately for seed region, microRNA (without seed region), pre-miRNA (without microRNA) and upstream and downstream pre-miRNA flanks. Note, that the MirSNP data (yellow bars) provide the densities of slightly different regions, namely, the entire pre-miRNA sequence and both 200bp pre-miRNA flanks. (B) SNP occurrence per pre-miRNA for disease or non-disease mirtrons and non-mirtrons. The disease pre-miRNAs are associated with at least one disease and were taken from [59, 60], SNPs were extracted from miRNASNP-2.0 [58].

The region regarding the canonical miRNAs can be divided into two parts of equal conservation: miRNA sequence and pre-miRNA (excluding the miRNA) with its flanks (figure 4A). This confirms only one relation in the previously found hierarchy of the SNP densities (seed < miRNA < pre-miRNA < flanks) [60, 72, 73]. In particular, the small difference between the seed and the rest of the miRNA (figure 4A, red bars) became even weaker (blue bars).

The miRNA sequence is saturated with functional sites (seed, additional binding site, etc.). Besides, miRNA duplex holds a most part of its pre-miRNA and thus carries the main burden of responsibility for forming the pre-miRNA hairpin. The rest of the pre-miRNA and its flanks also include a number of important regulatory elements (e.g. miRNA binding sites, UG, CNNC and other motifs [68, 75]), which are sparsly distributed over RNA sequence. This agrees with our observation that the pre-miRNAs and their flanks have greater SNP density than the miRNA sequences (figure 4A).

The other reason of hierarchy disappearing could be that the updates of miRBase and dbSNP cause the SNP density to increase throughout the regions and the difference between them to smooth off. For example, the dbSNP can rapidly accumulate rare SNPs and the miRBase – less conserved sequences. We consider this in the last section where we test the robustness of our results. As it turned out, the hierarchy of the SNP densities is being restored for common SNPs in the robust miRNAs.

In the mirtrons the SNPs take place more frequently than in other miRNAs and pre-miRNAs (figure 4A) in accordance with the fact that SNPs most often occur in introns [76]. Intronic SNPs can affect mRNA expression and splicing [77] and thus may influence the mirtron processing. In contrast with the non-mirtrons, SNPs proceed in the mirtron pre-miRNAs more frequently than in their flanks, in line with that the mirtron pre-miRNA flanks can often contain exons where SNPs rarely occur [76].

Lu et al. and later Han et al. explored the relation between miRNA conservation and diseases, and found out that SNPs occur less frequently in miRNAs associated with diseases than in non-associated ones [59, 60]. Using recent SNP data we re-calculated the SNP occurrence of non-mirtrons (figure 4B) and confirmed their observation that SNP-rare pre-miRNAs are often associated with a number of diseases while SNP-frequent ones are not [59, 60]. In contrast, the non-disease mirtron pre-miRNAs have lower number of the SNPs per sequence than the disease ones. Taking together the species specificity of mirtrons [45, 46], their increased SNP density (figure 4A) and their disease association, this suggests that mirtrons undergo positive selection while the most canonical miRNAs are under the negative one [78]. These also agrees with our observation that the mirtrons have wider distributions than the non-mirtrons (figures 2 and 4).

However, the results of such a simple SNP analysis as above should be taken with great caution: the carefully prepared samples, allele frequencies and other population genetic parameters are needed for a more profound analysis.

### Branchpoints in mirtrons and introns

Mirtrons are confirmed by splicing dependence of miRNA expression. And vice versa, the hairpin of future pre-miRNA may affect splicing, in particular, the branchpoint site recognition.

In a recent paper [61] authors extensivly studied the human, mouse and yeast branchpoints. Comparing their U2 basepairing modes we observe that the U2:mirtron model has been identified less frequently than the U2:intron model (table 1, Additional file 6): thus, more close inspection of mirtron splicing, in particular, the role of intronic hairpins immediately upstream the 3′ splice site, can shed a new light on splicing in general.

Using human and mouse data from Taggart et al. [61] we compare the branchpoint distribution of mirtrons and introns. Despite the distributions similarity, mirtron branchpoints are more often located in the expected region (10-40 nucleotides upstream from the 3′ splice site) than the introns what suggests the more frequent constitutive splicing of the mirtron precursors (figure 1, Additional file 6). Remarkably, the mirtron branchpoints are most probably located at 18-24nt away from 3′ splice site, i.e. near the miRNA end (figure 1, Additional file 6).

To characterize the branchpoint distribution more closely we analyzed its locations along pre-miRNA sequences and found that the branchpoint often appears in terminal loop or in 3' pre-miRNA strand (Figure 5A). In those cases when it was found in terminal loop, the pre-miRNA hairpin is most frequently as short as possible (figure 5B). When branchpoint is located in 3' strand, its site is more likely to be 6-8 nt away from the terminal loop overlapping with the Dicer cleavage site as it follows from the mirtron length distributions (figure 5B) and from direct calculation of distances between branchpoint and Dicer cleavage site (figure 5C). The latter site attracts a nearest loop (figure 2C) so we conclude that the mirtron pre-miRNA secondary structure tends not to shield the branchpoint (therefore not to block the U2:mirtron basepairing) but rather to fix branchpoint and 3′ splice site mutual disposition.

**Figure 5.**
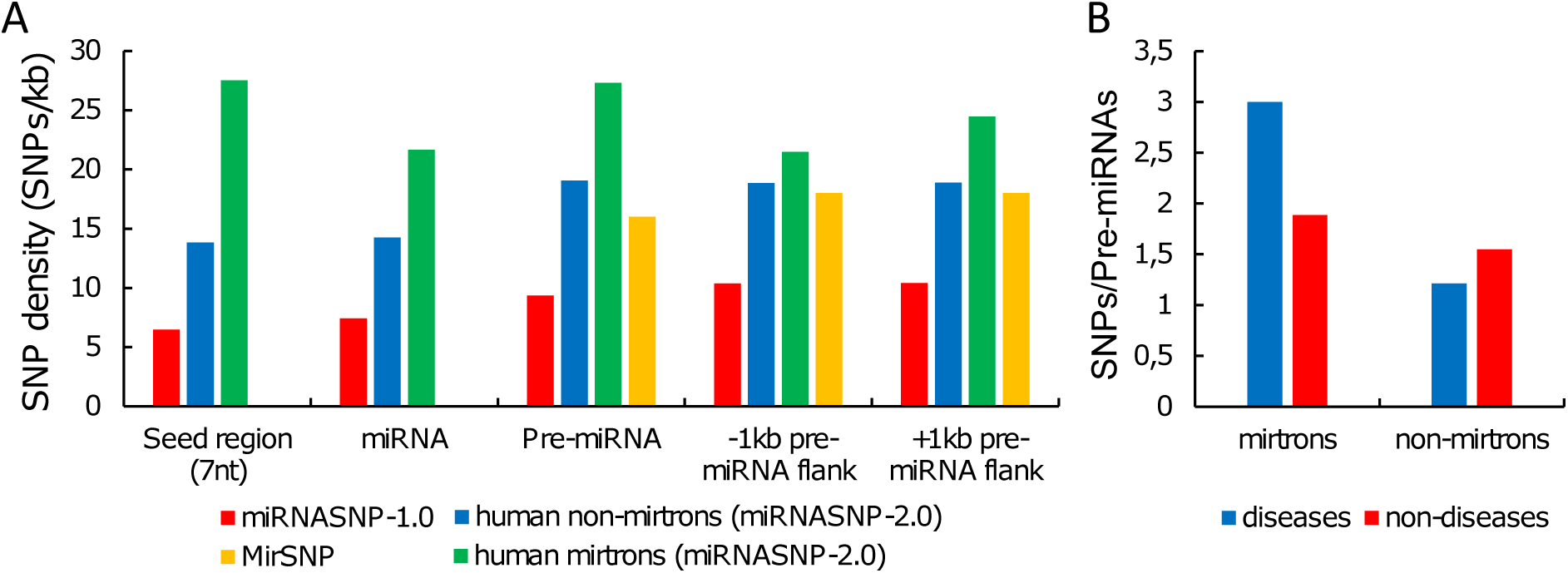
Distance from terminal loop to branchpoint (A) and to 3′ **pre-miRNA end (B)**. **Distance between branchpoint and Dicer cleavage site of 3**′ **miRNA (C).** Note that the 3′ miRNAs are mainly located near the 3′ splice site. The two separate points on the left figure show the branchpoint frequencies of the 5′ pre-miRNA strand (green) and of the terminal loop (red) as a whole. The blue curve displays the branchpoint distribution along the 3′ pre-miRNA strand. On the center figure shown are the distances of two mirtron groups: with branchpoint within the 3′ pre-miRNA strand (blue) and with branchpoint within the terminal loop (red). On the right figure considered are the distances between the 5′ end of the 3′ miRNA and the branchpoint within the 3′ pre-miRNA strand. Negative values correspond to the branchpoints into miRNA sequence.

### Checking the stability of the results

MiRBase is often criticized for including the transcriptional noise [57, 79–81] which sometimes is estimated to be as high as 2/3 of the annotated human miRNAs [57]. This criticism has motivated to filter the miRBase entries and to establish a new miRNA catalog (MirGeneDB database [57]).

To verify that our results are robust to the miRBase false positives, we repeated our computations for miRNAs whose identifiers are presented in MirGeneDB. The MirGeneDB authors rejected the most of the mirtrons, mainly by the improper mature/star offset and the undesirable heterogeneous processing. As a result, the MirGeneDB contains only 7 human and mouse mirtron entries (mmu-mir-1981, mmu-mir-3097, hsa-mir-3605, hsa-mir-3940, hsa-mir-4640, hsa-mir-5010, hsa-mir-6746). Therefore, we considered only the non-mirtrons.

We found that most of our results for non-mirtrons are stable against the miRBase reducing to robust miRNAs (Additional file 7). The main differences are the following. The miRNA regions became more contrasting, especially for the SNP densities. Generally, the SNP densities decreased up to 1.5-times and the difference in the densities of the pre-miRNA and its flanks re-appeared. After additionally removing the rare SNPs [82] the densities decreased much stronger and their hierarchy was completely restored (seed < miRNA < pre-miRNA < flanks, Additional file 7). The common SNPs are likely older, they have been subjected to selective forces over time [83] and produces the difference between seed and the rest of miRNA.

Most of variance, in particular within miRNAs, arises from rare variants most of which are either recently derived alleles or being selected against due to their deleterious nature [84]. This effect may be most pronounced in modern humans who live under relaxed selection and readily accumulate the deleterious rare alleles [85]. Mirtrons are more variable and likely carry more rare SNPs than the canonical miRNAs. Multiple rare SNPs often associate with complex diseases. Therefore we speculate that mirtrons are a rich source of disease-promoting variants (figure 4B).

## Conclusions

The pri-/pre-miRNA secondary structure plays an important role on each stage of biogenesis, in particular by positioning the miRNA excision sites. Drosha as well as Dicer can cleave imprecisely around the expected sites. For mirtrons splicing replaces the Drosha cleavage and manifests itself in the footprints of the secondary structure near the pre-miRNA base (dangling ends and less precise cleavage), while for the Dicer cleavage sites the characteristics of the canonical biogenesis matches the non-canonical ones. Both complexes recognize the inner/bulge-loop structure; therefore the loop-counting rule can help to predict not only Dicer cleavage site but also Drosha one. Dicer binds the pre-miRNA ends: as the result, the imprecise Drosha cleavage can induce Dicer error what appears in the dependence of the overhang lengths. To explain this interrelation of Dicer and Drosha precision we suggest the two-lever model of Dicer movements where the distance between RNase IIIA/IIIB cleavage sites fits the distance between pre-miRNA ends. In mirtron pre-miRNAs both ends are typically hanging and their distance varies widely thus increasing the Dicer cleavage imprecision. Also the mirtron hairpin brings together the splice sites for 60-80nt closer, exposes branchpoint site and adjusts it on the 3′ splice site. The mirtron structure appears to be well suited to the splicing and thus the mirtrons can evolve from the occasional hairpins (as readily as other miRNAs) in the immediate neighbourhood of the 3′ splice site. Also through the splicing mirtrons can acquire guanine at their ends what induces the Dicer impresicion.

The secondary structure of pri-/pre-miRNAs appears also in their evolution, in particular, in clear difference of mutation rates between single- and double-stranded positions: thus the secondary structure shapes the functional subdivision of the precursor (seed, additional binding site, inner and terminal loops, etc.). For the SNP density extracted from the last miRNA SNP database this division is reduced because of rare SNPs and non-robust miRNAs. In contrast to the canonical miRNAs the mirtrons exhibit higher SNP density and more SNPs per pre-miRNAs that are associated with diseases. This suggests that mirtrons unlike old canonical miRNAs are under positive selection and serve as an inherent source of silencing variability.

## List of abbreviations

UNF: unpaired nucleotide frequency

## Declarations

### Ethics approval and consent to participate

Not applicable.

### Consent for publication

Not applicable.

### Availability of data and material

All data analysed during this study are freely available from the sources in this published article.

### Authors' contributions

PSV performed bioinformatics calculations. IIT and PSV contributed to the interpretation of results, and were involved in manuscript editing. IIT designed and coordinated the study. All authors have read and approved the final manuscript.

## Acknowledgements

Not applicable.

## Additional files

Additional file 1. The particular examples of the 3′ end shifting inside miRNA. docx.

Additional file 2. Change of the pre-miRNA secondary structure reduces a pair of long overhangs to nearly canonical one. docx.

Additional file 3. The model of independent overhangs and ist results. docx.

Additional file 4. Overhang length dependence are robust against the overhang definition. docx.

Additional file 5. The scheme of the proposed 2-lever model for Dicer. png. Additional file 6. The branchpoint locations across mirtrons and introns. docx. Additional file 7. The results of the stability test using MiRGeneDB database. docx.

